# Algal growth at environmentally relevant concentrations of suspended solids: implications for microplastic hazard assessment

**DOI:** 10.1101/2020.04.11.036889

**Authors:** Elena Gorokhova, Karin Ek, Sophia Reichelt

## Abstract

Hazard assessment of microplastic is challenging because standard toxicity testing is mostly developed for soluble (at least partially) chemicals. Adverse effects can occur when test organisms are exposed to turbid environments with various particulate matter (PM), both natural, such as sediment, and anthropogenic, such as microplastic. It is, therefore, relevant to compare responses to PM exposure between the microplastic and other suspended solids present at ecologically relevant concentrations; this can be done by using reference materials when assessing hazard potential of microplastics. Here, we evaluated growth inhibition in unicellular alga *Raphidocelis subcapitata* exposed to different suspended solids (microplastic, kaolin, and cellulose; 10, 100 and 1000 mg/L) during 72 h; algae without added solids were used as a control. In addition, aggregate formation in the exposure systems was analyzed using particle size distribution data. At 10 and 100 mg/L, no adverse growth effects were observed in any treatments; moreover, algal growth was significantly stimulated in kaolin and cellulose treatments compared to the control. However, at 1000 mg/L, all tested materials exerted growth inhibition, with no significant differences among the treatments. The comparison among particle size distributions across the treatments showed that both PM concentration and size of the particle aggregates were significant growth predictors for all materials tested. Therefore, at high concentrations, both natural and anthropogenic materials have similar capacity to cause adverse effects in algal growth inhibition tests, which must be taken into account in hazard assessment of plastic litter.

## Introduction

The occurrence of plastic in the environment has raised concerns (Andrady and Neal, 2009). However, plastic litter, including microplastic, is a new field in environmental pollution research with much-unsettled methodology, and standard methods for hazard assessment are not yet available (Adam et al., 2019). Current test methods in ecotoxicology are initially intended for soluble (at least partially) chemicals, whereas testing particle suspensions, such as microplastic, requires different approaches (Paul-Pont et al., 2018). Therefore, the development of adequate methods for microplastic hazard assessment under reproducible settings is the key challenge of the plastic litter toxicology.

Despite environmental health concerns, only a few studies have shown consistent adverse effects of microplastics (Foley et al., 2018), and they are usually observed at the highest concentrations tested. As a rule, these concentrations exceed levels that are so far recognized as environmentally relevant (Lenz et al., 2016; Phuong et al., 2016). For such studies, a valid critique is a high probability that these effects are not necessarily specific to the microplastic but arise from the exposure to any suspended solids that have no nutrition value for animals and – in the case of photosynthetic microorganisms – a capacity to cause shading effects and sorb nutrients and microelements, thus decreasing their availability for the test algae (Ogonowski et al., 2018b). Hence, the existing protocols for hazard assessment do not explicitly address the effects of microplastic but those of particulate matter (PM) that change light and nutrient availability for the test organisms, such as algae and periphyton.

So far, most efforts have focused on animal responses to microplastic exposure (Foley et al., 2018; Ogonowski et al., 2018b), whereas far less research addressed possible effects of microplastic on primary producers (Yokota et al., 2017). The reported consequences of the interactions between primary producers and particles of fossil-based polymers in nano- and microparticle size range include alterations in algal photosynthesis (Bhattacharya et al., 2010), growth (Sjollema et al., 2016; Bergami et al., 2017a), and colony morphology (Yokota et al., 2017). These effects are important as they can propagate in the food web, affecting consumers and system productivity. However, the reported effects are not consistent across the studies, and no-effect reports are also published (Prata et al., 2019). What is also essential is that similar effects on the same endpoints can be induced by natural suspended solids, such as clay and sediment (Bilotta and Brazier, 2008; Chapman et al., 2017). This is not particularly surprising because the negative effects of fine sediments on algae are well-known from the field studies (Cahoon et al., 1999). It is, therefore, relevant to compare responses to particle exposure between the microplastic and other suspended solids present at ecologically relevant concentrations in the environment.

Microalgae have a long history of use in ecological and ecotoxicological assessments because of their high sensitivity towards various stressors, including environmental pollutants. In ecotoxicology, the standard algal growth inhibition test (OECD, 2006) validated for several freshwater and marine species is widely used for hazard evaluation. When conducting this test with various effluents, it is recommended to measure total suspended solids and total settled solids, because removal of these fractions of the effluent could influence the growth inhibition and thus toxicity estimates. The mechanisms of the growth inhibition in the presence of suspended solids, including microplastic, may vary, including shading effects and trapping of microalgae in the aggregates and subsequent inhibition of photosynthesis (He et al., 2017; Mao et al., 2018). Recently, the latter effects have been suggested to occur in the algae exposed to microplastic; moreover, increased production of organic carbon and its aggregation into gel particulates have been demonstrated in mesocosms amended with polystyrene microbeads (Galgani et al., 2019). It is, therefore, crucial to understand whether algae-plastic aggregates have a higher capacity to inhibit growth compared to the aggregates with natural solids.

Also, as a consequence of degradation processes, the plastic particles in the environment undergo weathering that changes their physicochemical properties and, possibly, hazard potential. Various processes, such as physical stress, UV-radiation, shifting temperatures, salinity and oxidization contribute to weathering resulting in changes of the surface properties and morphology (Jahnke et al., 2017). Therefore, it is questionable whether the common use of uniform spherical MP in experiments is justifiable from ecological and ecotoxicological viewpoints. Hence, using aged polymer fragments to better mimic the environmentally realistic exposure scenario has been advocated (Ogonowski et al., 2018b). It is, however, only a very few studies that have evaluated the weathering effect in MP toxicity tests (Bråte et al., 2018; Fu et al., 2019), and found that aging of MP (e.g., PVC) would pose stronger inhibitory effects on microalgae (Fu et al., 2019).

The objectives of this study were to (1) compare effects of fossil-based polymers and natural (kaolin and cellulose) particles on the growth performance in the standard algal growth inhibition test (OECD, 2006); (2) compare algal growth response to suspended solids between weathered and virgin microplastic; and (3) evaluate relationships between particle aggregation in the experimental system and growth rate of the algae. The present study does not aim to generate a robust test design for suspended solids in general or for microplastic specifically but rather is intended to generate debate on the methodology of microplastic hazard assessment and possibly influence further research efforts toward this goal.

## Material and Methods

### Test organism

The freshwater unicellular green alga *Raphidocelis subcapitata* (Korshikov) Nygaard, Komárek, J. Kristiansen & O. M. Skulberg, 1987, formerly *Pseudokirchneriella subcapitata* and *Selenastrum capricornutum*, is a standard test organism in ecotoxicology (OECD, 2006). It is a fast-growing species, sensitive to light and nutrients, and thus particularly well suited to evaluate stress effects on algal growth and production (Gonçalves et al., 2016). Algal culture for inoculation was grown for one week in Z8 media, with shaking (100–125 rpm) at room temperature and the illumination of ~40 μ E·m^−2^·s^−1^. Algal concentrations were determined by *in vivo* fluorescence using a 10 AU™ Field Fluorometer (Turner Designs, Sunnyvale, California, US).

### Chemicals, reference, and test materials

Kaolin (Sigma-Aldrich, K7375) and native fibrous cellulose (Macherey-Nagel, MN 301) were used as the reference materials. Kaolin contains mainly the clay mineral kaolinite, a hydrous aluminosilicate, whereas cellulose is the primary substance in the walls of plant cells. Both materials occur globally in suspended particulates and have been used as reference material when assessing microplastic effects (Gerdes et al., 2019) and as a test material when assessing the effects of total suspended solids (Robinson et al., 2009; Ogonowski et al., 2018a). As a test microplastic, we used polyethylene terephthalate (PET, Goodfellow GmbH, product number ES306312) mixed with Milli-Q water passed through a 40-μm sieve to produce a size fraction similar to that of kaolin and cellulose. To investigate whether weathering of microplastic affects aggregating behavior and effects on the algae, we used both virgin and weathered PET particles, denoted as PET and PET_w_, respectively. For testing, PET_w_ was prepared by UV exposure of the milled PET for 28 days (Oelschlägel et al., 2018) and size-fractionated using the same technique as PET (Text S1; Supporting Information). Stocks of both reference and test materials contained 0.01% v/v of a non-ionic surfactant (Tween-80, Sigma-Aldrich); it was also added to the control media at the final concentration of 0.0001%. In the experimental treatments, the concentration of Tween-80 never exceeded 0.0001%, which is well below the levels regarded as non-toxic for green algae (Ma et al., 2004).

In the stock suspensions, both PET materials (median size ~11 and ~12 μm for virgin and weathered PET, respectively) and kaolin (~9 μm) showed unimodal size distributions, whereas cellulose (~16 μm) had polymodal distribution and broader size range. See Text 1, Table S1, and Fig. S1 for the description of the particle preparation and size distributions of the stock suspensions used to prepare exposure media at different nominal concentrations of suspended solids.

Standard growth media (Z8) contained all nutrients at surplus for the entire growth period of the algae in the incubation system: NaNO_3_ (6 mM), CaCl_2_·2H_2_O (73 mg L^−1^), MgSO_3_·7H_2_O (25 mg L^−1^), K_2_HPO_4_ (31 mg L^−1^), Na_2_CO_3_ (21 mg L^−1^), FeCl_3_·6H_2_O (28 mg/L) 0.001 M HCl, ZnSO_4_·7H_2_O (5 μ g/L), MnCl_2_·_4_H_2_O (10 μ g/L), H_3_BO_3_ (5 μg L^−1^), CuSO_4_·5H_2_O (0.5 μg L^−1^), CoCl_2_·6H_2_O (2 μg L^−1^), NaMoO_4_·2H_2_O (1.5 μg L^−1^), VOSO_4_·2H_2_O (0.5 μg L^−1^), Na_2_SeO_4_·10H_2_O (0.5 μg L^−1^), pH adjusted to 6.6–7.0. The chemicals were purchased from Sigma-Aldrich (Germany).

### Experimental setup

In total, five experimental media with different suspended solids were prepared by mixing stock suspensions of the test material and used as treatments with nominal concentrations of suspended solids 10, 100, and 1000 mg L^−1^ (Table 1). This design resulted in twelve different combinations of material × concentration factors for algal exposure and a single control with no added suspended solids; five replicates were used for each of these incubations. Also, particle blanks were used for each material × concentration combination (Table 1) to control for contamination with microorganisms (algae and bacteria) and any background fluorescence induced by the test and reference materials under experimental conditions; all particle blanks were run in triplicates. When measuring fluorescence in the treatment tubes, the mean value of the particle blanks was subtracted from each replicate for the respective material × concentration combination.

**Table 1.** Experimental setup for growth inhibition assay with *R. subcapitata*. Algae were exposed to test microplastics, virgin and weathered polyethylene terephthalate (PET and PET_w_, respectively), and reference materials (kaolin and cellulose) at concentrations 10, 100 and 1000 mg L^−1^; control (no added suspended solids) and particle blanks (no added algae) were included. Standard growth media (Z8) contained all nutrients at surplus (Supplementary Information, Text S1).

| Treatment | Experimental media | Algae | Concentration of SS, mg L<sup>-1</sup> |
| --- | --- | --- | --- |
| Control | Z8 | yes | 0 |
| Kaolin | Z8 + Kaolin | yes | 10, 100, 1000 |
| Kaolin blank | Z8 + Kaolin | no | 10, 100, 1000 |
| Cellulose | Z8 + Cellulose | yes | 10, 100, 1000 |
| Cellulose blank | Z8 + Cellulose | no | 10, 100, 1000 |
| PET | Z8 + PET | yes | 10, 100, 1000 |
| PET blank | Z8 + PET | no | 10, 100, 1000 |
| PET<sub>w</sub> | Z8 + PET<sub>w</sub> | yes | 10, 100, 1000 |
| PET<sub>w</sub> blank | Z8 + PET<sub>w</sub> | no | 10, 100, 1000 |

A standard growth inhibition assay with *R. subcapitata* was conducted (OECD, 2006) with some modifications. The assay is based on the measurement of *in vivo* chlorophyll *a* (Chl *a*) that exhibits endogenous red fluorescence (autofluorescence). The quantification of chlorophyll fluorescence is useful for monitoring photosynthetic capacity and detection of stimulating or inhibiting effects. The algae were harvested from the culture and diluted with Z8 media to 10^5^ cell mL^−1^ and then mixed with the experimental media (Table 1) to the final cell density of 5 × 10^3^ cells mL^−1^. These suspensions were transferred to 10-mL glass tubes filled to the top and sealed to avoid air bubbles. All tubes were mounted on a plankton wheel rotating at 0.5 rpm and incubated at room temperature and fluorescent light at 140 ± 10 μ E·m^−2^ s^−1^. Light intensity was measured in the exposure area using photosynthetically active radiation sensor Quantum (Skye, UK). The algae were allowed to grow throughout the lag phase and exponential growth for 72 h, and fluorescence was measured at time points 0, 24, 48, and 72 h. The mean fluorescence of the particle blanks (Table 1) was subtracted from each replicate in the respective treatments. Over the range of cell densities used in this experiment, the fluorescence was linearly related to the cell number as established by the comparison between the fluorescence measurements and cell counts with a hemocytometer. Upon termination of the experiment, samples of the suspended matter were collected from each replicate and used for particle size distribution analysis in order to establish relationships between the growth parameters and particle aggregation.

### Growth analysis

The algal growth response was inferred from the chlorophyll *a* fluorescence every 24 h over 72 h. Growth kinetics of the algae was examined using time-specific measurements fitted to an exponential growth curve with lag phase (Baranyi and Roberts, 1994); see Supporting Information Text S2 for the calculation details. With DMFit software (www.combase.cc), the lag phase duration (λ, h) and maximal growth during exponential phase (μ, d^−1^) of the growth curves were estimated for each treatment. The lag phase duration reveals how fast test organisms acclimate to specific conditions, while the growth rate in the exponential phase indicates proliferation in the adapted population. Model fit was evaluated by the coefficient of determination (R^2^) and performance by the root mean square error (RMSE). An additional parameter, the change in fluorescence intensity between the observation points, was used to calculate the area under the curve (AUC). This endpoint is commonly used in algal growth inhibition assay as it integrates the change in photosynthetically active biomass during both the lag and growth phases representing the productivity of the population (Vaas et al., 2012).

### Particle distribution analysis

The particle size distribution (PSD) was measured with a Spectrex laser particle counter (Spectrex, model PC-2000, Redwood City, USA). The size spectra were determined for 1 – 100 μm range (Supporting Information Text S1) and processed with GRADISTAT program, version 8.0 (Blott and Pye, 2001) according to the method by Folk and Ward (Folk and Ward, 1957). Using PSD data, mean particle size, sample sorting (variance), skewness, kurtosis, median particle size (D_50_), and two points which describe the coarsest and the finest parts of the distribution (D_90_ and D_10_, respectively), and D_90_ – D_10_ range were obtained. These parameters are commonly used in concert for PSD characterization in a powder or granular material, such as soil and sediment (Blott and Pye, 2001).

### Statistics

We used two-way ANOVA to evaluate the effects of suspended solids concentration (SS, mg L^−1^) and test material on the algal growth parameters (λ, μ, and AUC) in different treatments. When the *material × SS* interaction was significant, the effect of *material* was evaluated for each level of concentration using Fisher LSD test for multiple comparisons. The data were not transformed as the model residuals were normally distributed. Results are expressed as mean value ± standard deviation (SD). Within-treatment variability was accessed as the coefficient of variation (CV%).

The PSD data from GRADISTAT for all treatments were first examined by Principal Components Analysis (PCA) to (1) explore the overall differences between the treatments with and without algae, (2) examine the relationships between PSD characteristics, and (3) identify the leading components. Then, we used Generalized Linear Model (GLM, log-link, normal error structure) analysis to test for significant effects of the algae in the system (present/absent) and SS concentration (0 to 1000 mg/L) for each material (kaolin, cellulose, PET and PET_w_) as predictors and each PSD metric (mean particle size, variance, skewness, kurtosis, D_50_, D_90_, D_10_, and D_10_ – D_90_) as a dependent variable.

Partial least square regression (PLSR) (JMP^®^, Version 14.0. SAS Institute Inc., Cary, NC, 1989-2019) was used to study the relationships between the exposure variables (PSD metrics and SS concentration; X variables) and the growth parameters (λ, μ, and AUC; Y variables); see Text S3 for details. The analysis was carried out for each material as well as for the pooled data set using the non-linear iterative partial least squares (NIPALS) algorithm; the data were centered and scaled. For cross-validation and determination of the optimal number of latent variables, we applied the leave-one-out method and predicted residual error sum of squares (PRESS) statistic as implemented in the PLS platform of the software to separate terms that do not make important contribution to the dimensionality reduction involved in PLSR (Variable Importance in Projection; VIP < 0.8) and those that might (VIP ≥ 0.8). For each X variable, VIP value was used to assess its importance in the determination of the PLSR projection model (Wold et al., 2001). Also, the percentage of variation explained for X and Y variables, and the contribution of each of the important factors were assessed. To give another view on the relationships between the X- and the Y-variables and the PLSR factors, we calculated and plotted loadings.

## Results

### Algal growth response to suspended solids

Positive growth and growth models with high R^2^ values (> 0.97 in all cases) were observed in all treatments during the exposure, albeit with different growth trajectories (Fig. S1). There were significant effects of exposure on all growth parameters (λ, μ, and AUC), both positive and negative compared to the particle-free control (Table 1); these effects were pronounced in the algae exposed to the microplastic as well as in the reference materials (Fig. 1).

**Fig. 1.**
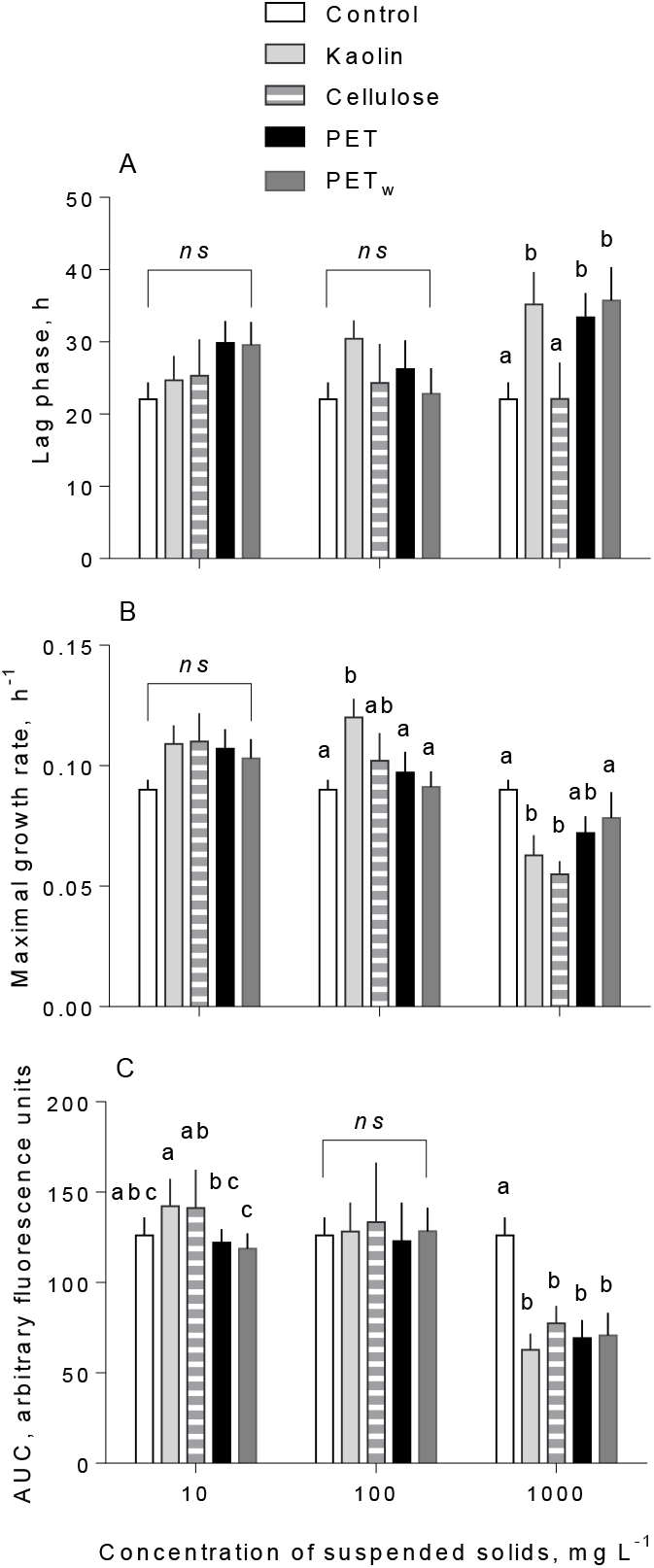
Kinetic parameters (mean and SD; *n* = 5) of growth responses in *R. subcapitata* exposed to microplastics and reference materials (A) Lag phase (λ) preceding the active growth, (B) maximal growth rate (μ) estimated for the exponential period, and (C) area under curve (AUC) representing algal production during the observation period. Nonmatching letters indicate a significant treatment difference at p < 0.05 evaluated by the Fisher LSD multiple comparisons test; *ns* denotes lack of significant differences between the materials within the test concentration of suspended solids; see Table S2 for the details.

Both λ and μ values varied among the treatments, with particular deviations observed at the highest concentration (1000 mg L^−1^). The lag phase was significantly higher in the algae exposed to 1000 mg L^−1^ for all materials tested, except cellulose, compared to that in control (Fig. 1A). Significant deviations from the control were observed for the maximal growth rate in the algae exposed to kaolin, with a significant increase at 100 mg L^−1^ and a decrease at 1000 mg L^−1^, whereas there were no significant effects for the other materials (Fig. 1B). Most pronounced changes were observed for the AUC values indicating a significantly lower production in PET_w_ treatment compared to the reference materials already at 10 mg L^−1^ and a decrease in all tested materials at 1000 mg L^−1^ (Fig. 1C). Notably, the within-treatment variability for AUC decreased significantly with time and increased with concentration (Table S1; Fig. 2). At each test concentration, the *material* × *time* interaction effect on CV% was not significant (Table S1), indicating that in all treatments, *material* effect was persistent.

**Fig. 2.**
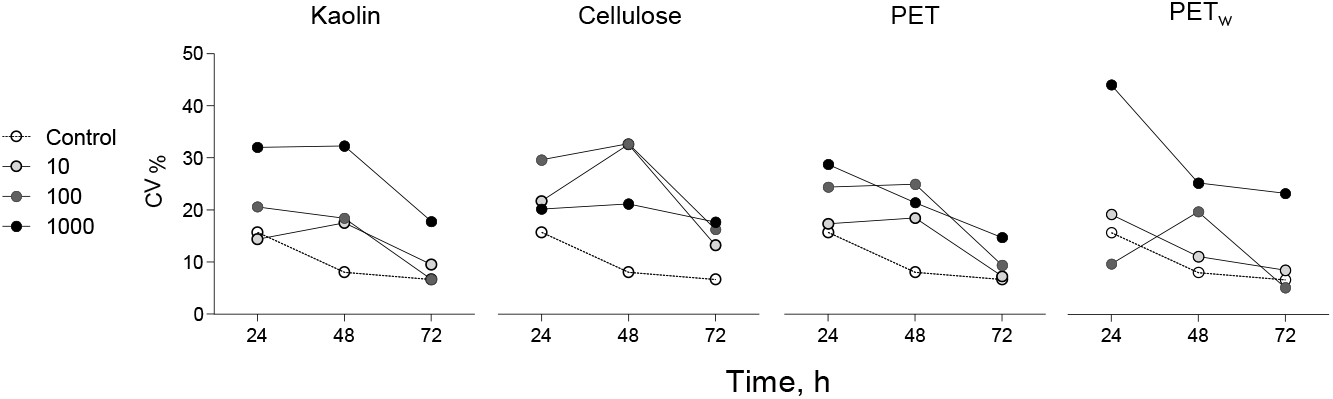
Change in coefficient of variation (CV%) of AUC values for growth curves of *R. subcapitata* exposed to microplastics and reference materials over the course of the experiment (72 h). The AUC values were grouped by test material (kaolin, cellulose, PET and PET_w_) and test concentration (10, 100 and 1000 mg L^−1^; *n* = 5); control values are shown on each plot for comparison. See Table X for statistical evaluation.

There was an overall positive relationship between the maximal growth rate and lag phase duration across the treatments at concentrations 10 and 100 mg L^−1^ (Table 2). However, algae exposed to 1000 mg L^−1^ were deviating significantly from this relationship (Fig. 3), because of the significantly lower μ values in all treatments for this concentration.

**Table 2.** Outcome of two-way ANOVAs testing effects of suspended solids concentration (SS) and test material on the growth parameters (λ, μ, and AUC). When the *Material × SS* interaction was significant, the material effect was evaluated for each level of concentration using Fisher LSD test for multiple comparisons; the *post-hoc* results are presented in Fig. 1 and Table S2, Supplementary Information.

|  | SS | DF | MS | F | *p* value |
| --- | --- | --- | --- | --- | --- |
| **Lag phase, $\lambda$** |  |  |  |  |  |
| Interaction | 573.0 | 8 | 71.62 | 0.9955 | 0.4489 |
| SS | 277.3 | 2 | 138.6 | 1.927 | 0.1545 |
| Material | 856.7 | 4 | 214.2 | 2.977 | **0.0261** |
| Residual | 4317 | 60 | 71.94 |  |  |
| **Max growth rate, $\mu$** |  |  |  |  |  |
| Interaction | 0.007348 | 8 | 0.0009185 | 2.877 | **0.0089** |
| SS | 0.01549 | 2 | 0.007747 | 24.27 | **< 0.0001** |
| Material | 0.0006326 | 4 | 0.0001581 | 0.4953 | 0.7392 |
| Residual | 0.01916 | 60 | 0.0003193 |  |  |
| **Area under curve, AUC** |  |  |  |  |  |
| Interaction | 11150 | 8 | 1394 | 5.976 | **< 0.0001** |
| SS | 37995 | 2 | 18998 | 81.46 | **< 0.0001** |
| Material | 4636 | 4 | 1159 | 4.970 | **0.0016** |
| Residual | 13992 | 60 | 233.2 |  |  |

**Fig. 3.**
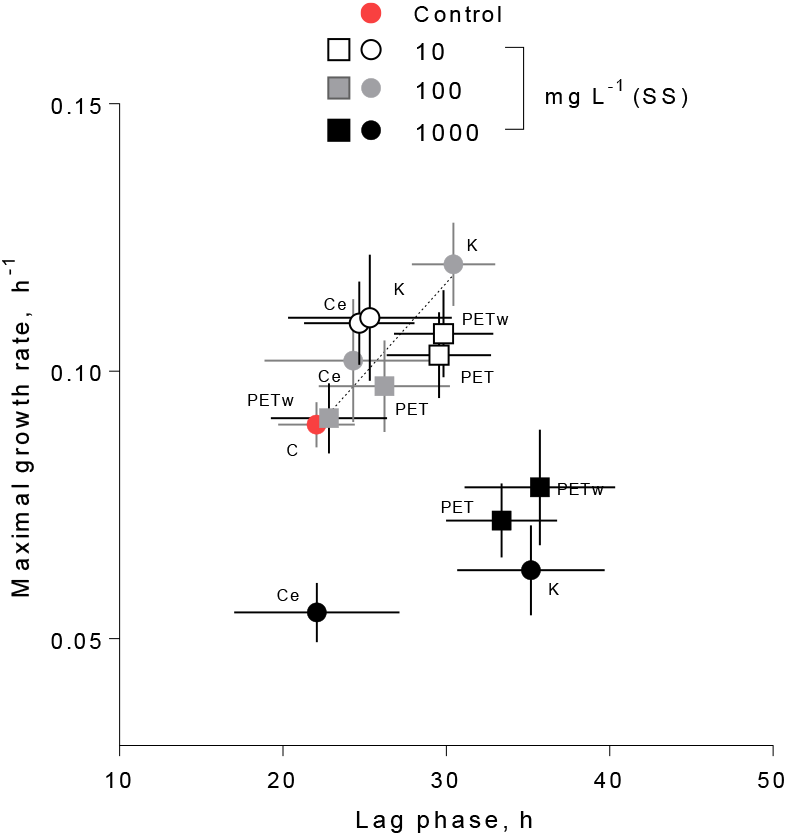
Relationship between maximal growth rate (μ, d^−1^) and duration of the lag phase (λ, h) in *R. subcapitata* exposed to microplastic and reference materials at concentrations of 10, 100 and 1000 mg L^−1^. No suspended matter was added to the medium in the control. Data are shown as means and SD (*n* = 5).

### Particle aggregation analysis

All types of PM aggregated during the exposure compared to the respective stocks. Moreover, they aggregated significantly more in the presence of algae compared to the particle controls, as evidenced by the particle distribution (Table 3; Fig. 5, Fig. S3). Principal components analysis (PCA) based on the PSD characteristics for each material and treatment indicated that two principal components exceeded 5% of the total explained variance (Fig. 4). Moreover, 83-94% and 6-15% of the accumulated dispersion were represented by PC1 by PC2, respectively, while only PC1 passed the broken-stick test (Fig. S4). The PCA biplot demonstrated separation between the treatments with and without algae for all four materials, with particularly strong separation observed for cellulose and PET_w_, whereas some overlap occurred for kaolin and PET treatments (Fig. 4). The loadings on PC1 indicate the importance of the particle size metrics (mean and median particle sizes, D_10_, D_90_, and D_90_-D_10_), whereas variance, skewness, and kurtosis were not influential for the discrimination between the treatments with and without algae (Fig. S5).

**Table 3.** Outcome of Generalized Linear Models (GLMs) testing effects of the algae (*Algae*) and suspended solids concentration (*SS*, mg L^−1^) on the PSD characteristics for each reference and test material. See Fig. X for the direction of the effects; *: *p* < 0.05; **:*p* < 0.01; ***: *p* < 0.001.

| Material | PSD characteristics | Wald Stat. for |  |  |
| --- | --- | --- | --- | --- |
|  |  | Main effects |  | Interaction |
|  |  | Algae | SS | *Algae* × *SS* |
| Cellulose | Mean size | 257.65*** | 461.64*** | 2.38 |
|  | Variance | 10.18** | 2.78 | 0.17 |
|  | Skewness | 7.96** | 4.30* | 0.70 |
|  | Kurtosis | 10.12** | 6.02* | 26.59*** |
|  | D<sub>10</sub> | 33.06*** | 103.71*** | 0.04 |
|  | D<sub>50</sub> | 217.46*** | 213.02*** | 9.04** |
|  | D<sub>90</sub> | 52.87*** | 343.19*** | 11.84** |
|  | D<sub>90</sub> – D<sub>10</sub> | 1.64 | 24.61*** | 7.30* |
| Kaolin | Mean size | 7.54* | 0.39 | 0.57 |
|  | Variance | 4.20* | 20.18*** | 5.38* |
|  | Skewness | 3.64 | 22.85*** | 7.66* |
|  | Kurtosis | 5.71* | 13.57*** | 23.98*** |
|  | D<sub>10</sub> | 11.79*** | 1.42 | 1.78 |
|  | D<sub>50</sub> | 8.05** | 0.07 | 0.82 |
|  | D<sub>90</sub> | 3.59 | 0.49 | 0.17 |
|  | D<sub>90</sub> – D<sub>10</sub> | 0.04 | 3.19 | 2.16 |
| PET | Mean size | 13.72** | 0.40 | 3.49 |
|  | Variance | 5.38* | 1.44 | 2.02 |
|  | Skewness | 5.02* | 1.42 | 2.72 |
|  | Kurtosis | 0.22 | 1.49 | 0.04 |
|  | D<sub>10</sub> | 0.01 | 0.55 | 11.77** |
|  | D<sub>50</sub> | 10.95** | 0.01 | 3.20 |
|  | D<sub>90</sub> | 9.02** | 3.63 | 5.69* |
|  | D<sub>90</sub> – D<sub>10</sub> | 7.55* | 3.64 | 1.97 |
| PET<sub>w</sub> | Mean size | 360.68*** | 265.31*** | 56.20*** |
|  | Variance | 1.41 | 4.16* | 13.91** |
|  | Skewness | 0.50 | 1.20 | 10.86** |
|  | Kurtosis | 0.67 | 2.73 | 5.89* |
|  | D<sub>10</sub> | 266.60*** | 59.24*** | 41.71*** |
|  | D<sub>50</sub> | 195.99*** | 99.81*** | 28.67*** |
|  | D<sub>90</sub> | 68.08*** | 103.95*** | 0.15 |
|  | D<sub>90</sub> – D<sub>10</sub> | 16.08*** | 56.64*** | 1.06 |

**Fig. 4.**
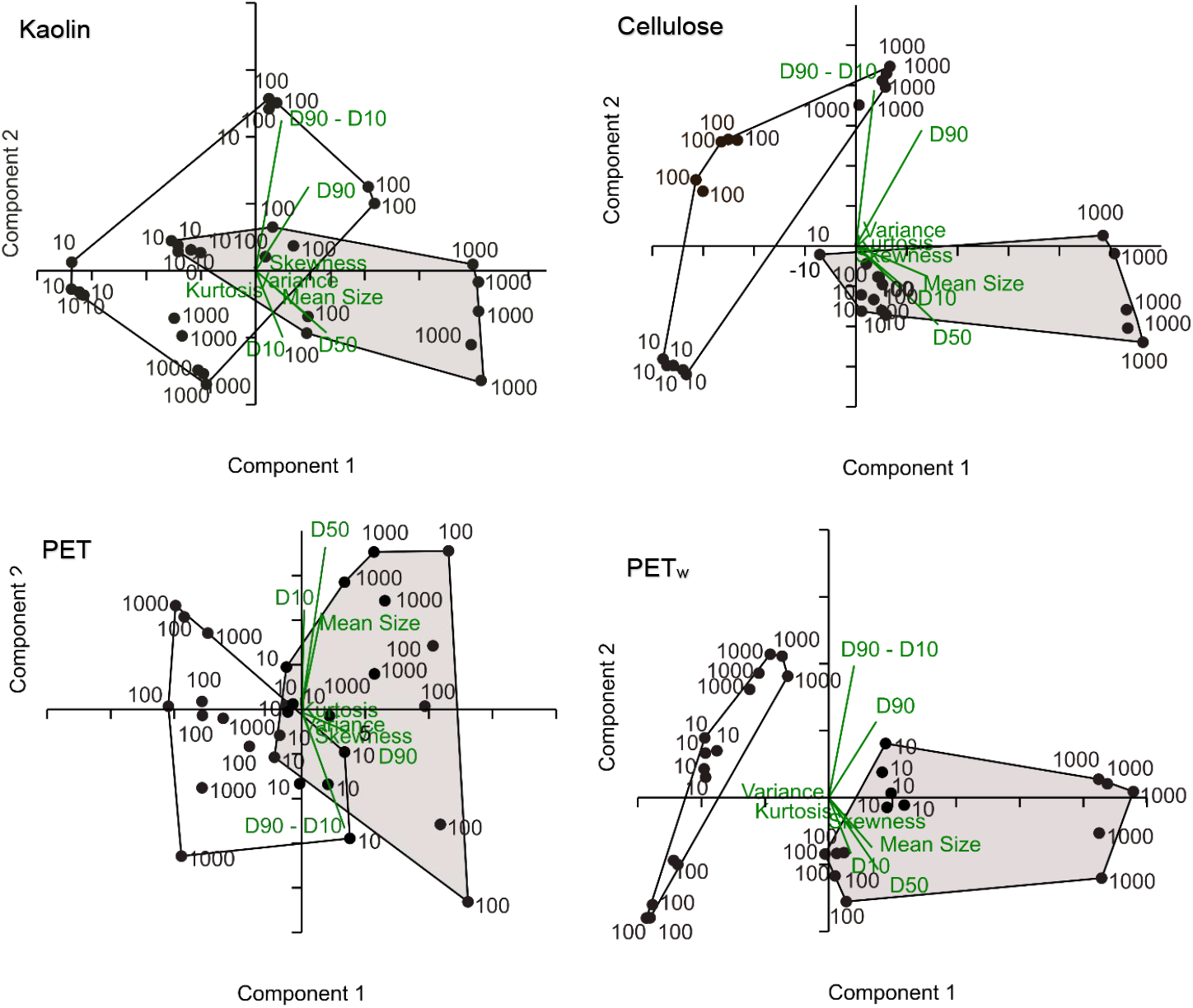
PCA biplot for the PSD characteristics obtained with GRADISTAT for the reference (kaolin and cellulose) and test (PET and PET_w_) materials in the treatments with algae (shaded convex hulls) and without algae (open convex hulls). The numbers indicate test concentrations (10, 100 and 1000 mg L^−1^; *n* = 5). The measurement vectors are shown in green.

**Fig. 5.**
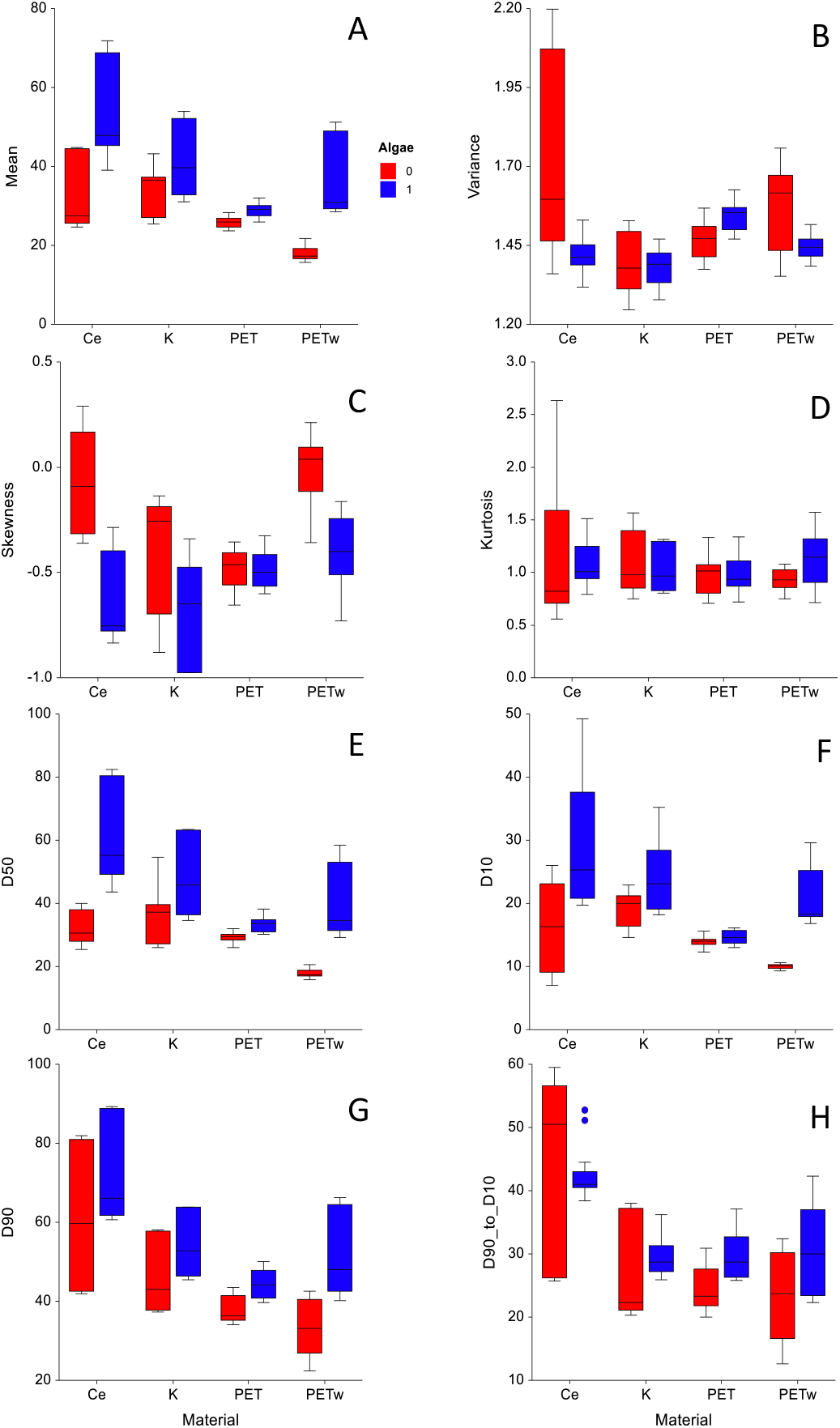
Different characteristics of particle size distribution (PSD) for the reference (kaolin and cellulose) and test (PET and PET_w_) materials determined at the termination of the incubation (72-h) with *R. subcapitata (Algae* = 1) and those from the particle controls incubated under the same conditions but without the algae (*Algae* = 0). The PSD parameters were as follows: (A) mean size, (B) variance (defined as sorting), (C) skewness, (D) kurtosis, (E) D_50_, (F) D_10_, (G) D_90_, and (H) D_90_-D_10_. Data are shown as box-and-whiskers (median value, 75% and 25%-quartiles, mix and max values; *n* = 5 in all cases).

### Effects of particle aggregation on algal growth

Significant PLSR models for growth parameters (Y variables) on PSD and SS concentration in the exposure systems (X variables) were obtained for all materials tested (Table 4; Fig. S6). The predictive capacity for all models was high, with Q^2^ varying from 74 to 96% (Table 4). In these models, 90% to 99% of the variance in the X-parameters explained 66% to 71% of the variance of the Y-variables (Table 4). Compared to the Lag phase, maximal growth rate and AUC were better predicted by the X-variables, particularly in the algae exposed to kaolin and PET (Fig. S7). In all models, SS concentration, mean, and median particle sizes were identified as significant growth predictors (Table 4); other PSD characteristics in different combinations for different materials were also important (Fig. S6).

**Table 4.** Significant PLSR models (*p* < 0.05) with growth parameters as dependent variables (all models) and PSD characteristics of the suspended solids and their concentration (SS) as explanatory variables for algae exposed to each reference and test material. The explanatory variables are listed in the order of importance based on their VIP score in the final model. All models comply with the quality criteria of (Lundstedt et al., 1998).

| Response variables | Explanatory variables | Number of factors | R<sup>2</sup>X | R<sup>2</sup>X<sub>cum</sub> | R<sup>2</sup>Y | R<sup>2</sup>Y<sub>cum</sub> | Q<sup>2</sup> | Q<sup>2</sup><sub>cum</sub> | Root mean PRESS |
| --- | --- | --- | --- | --- | --- | --- | --- | --- | --- |
| Max growth rate, Lag phase, AUC | Kaolin |  |  |  |  |  |  |  |  |
|  | D<sub>90</sub>-D<sub>10</sub>, Mean size, SS, D<sub>50</sub>, Skewness, D<sub>90</sub>, D<sub>10</sub> | 1 | 0.755 | 0.755 | 0.512 | 0.512 | 0.627 | 0.627 | 0.817 |
|  |  | 2 | 0.126 | 0.881 | 0.127 | 0.639 | 0.618 | 0.857 | 0.817 |
|  |  | 3 | 0.108 | 0.989 | 0.055 | 0.696 | 0.638 | 0.948 | 0.780 |
|  | Cellulose |  |  |  |  |  |  |  |  |
|  | Variance, Mean size, D<sub>50</sub>, D<sub>90</sub>-D<sub>10</sub>, Kurtosis, SS, D<sub>90</sub>, D<sub>10</sub> | 1 | 0.762 | 0.762 | 0.486 | 0.487 | 0.512 | 0.513 | 0.823 |
|  |  | 2 | 0.134 | 0.897 | 0.221 | 0.709 | 0.467 | 0.740 | 0.706 |
|  | PET |  |  |  |  |  |  |  |  |
|  | SS, D<sub>50</sub>, Mean size | 1 | 0.603 | 0.603 | 0.572 | 0.572 | 0.556 | 0.556 | 0.764 |
|  |  | 2 | 0.375 | 0.979 | 0.083 | 0.655 | 0.782 | 0.903 | 0.714 |
|  | PET<sub>w</sub> |  |  |  |  |  |  |  |  |
|  | Variance, D<sub>50</sub>, Mean size, Skewness, D<sub>90</sub>-D<sub>10</sub>, D<sub>90</sub>, D<sub>10</sub>, SS | 1 | 0.780 | 0.780 | 0.557 | 0.557 | 0.808 | 0.808 | 0.779 |
|  |  | 2 | 0.141 | 0.920 | 0.136 | 0.694 | 0.789 | 0.960 | 0.727 |

For all materials, the effects of particle size (mean size and D_50_) were adverse for the overall growth performance (Fig. S6). For kaolin, the SS concentration effect on the maximal growth and AUC values was weakly positive, whereas, for all other materials, it was negative, with the most substantial effects observed in the algae exposed to virgin PET. Other significant variables in the PLSR models were variance and skewness that were positive predictors in Cellulose and PET_w_ treatments, whereas the range interval D_90_-D_10_ was an important negative predictor for kaolin and PET_w_ (Fig. S6).

## Discussion

A step towards quantifying hazardous properties of synthetic polymers is to move the field of plastic litter toxicology beyond its current exploratory state to experimental designs that provide delineation of microplastic effects from those of suspended solids. Parallels can be drawn across particles and materials, regardless of whether they are natural solids, engineered nanomaterials, or plastics. On the other hand, it is essential to understand where the similarities end, in order to avoid redundant testing and use of inappropriate test methods.

Using a combination of the standard growth inhibition test and particle size distribution analysis, we found that all materials tested inhibited growth at the highest concentration (1000 mg/L) but not at the lower concentrations. Similar results were obtained for a green alga *Scenedesmus obliquus* exposed to polystyrene (PS; 0.07 μm particle size) by Besseling and co-workers (Besseling et al., 2014), who reported ~3% inhibition rate at 1000 mg/L and non-significant effects at lower concentrations. However, for another green alga *Dunaliella tertiolecta*, even lower concentrations of similar test material (PS; particle size 0.05 μm, 250 mg/L) caused a severe growth inhibition (57%) (Sjollema et al., 2016). However, the effect mechanisms for nanoparticles are likely to be different from the micrometer-sized particles, as shown for nanoparticle-cell interactions that involve extracellular proteins in the algal cell membrane with downstream effects on nutrient acquisition (Yue et al., 2017) or changes in colloidal stability affecting bio-physical interactions with cell walls (Gonzalo et al., 2014). In line with this, Nolte and co-workers using *Raphidocelis subcapitata*, i.e., the same green alga as in the present study, exposed to functionalized PS (0.11 μm) found 20-50% growth inhibition at concentrations 10 to 100 mg/L (Nolte et al., 2017). Therefore, it is most likely the particle size and not the material itself that would govern the algal responses in growth inhibition assays.

The observed responses may also vary during the exposure, thus setting specific requirements for the test duration. For instance, in a green alga *Chlorella pyrenoidosa*, a reduction in growth was observed during the lag phase, with some inhibition in the exponential phase, but during the stationary phase this trend was reversed due to a compensatory growth that exceeded the values observed in control (Mao et al., 2018). Such biphasic responses indicate that even when growth inhibition occurs, the algal populations were able to adapt and sustain productivity. We found that at 10 and 100 mg/L there was a significant positive correlation between the lag phase and the maximal growth rate during the exponential phase suggesting a compensatory growth due to adaptation during the exposure in the algal populations. We also observed differential responses for different growth parameters, with the lag phase being the least affected by the test material (Fig. 1A) and the total productivity represented by AUC values showing the most deviations from the controls (Fig. 1C). Notably, while the lag phase in the lowest concentration was prolonged (albeit, not significantly), the compensatory growth in 10 and 100 mg/L treatments was manifested as higher maximal growth during the exponential growth phase and the resulting total production that was significantly higher in the treatments with reference materials at 10 mg/L but not the polymers (Fig. 1). At 100 mg/L, although a significant compensatory growth was observed in kaolin (Fig. 1B), it was not sufficient to support the same production as in control (Fig. 1C), and at 1000 mg/L, the significantly prolonged lag phase and lowered maximal growth rate resulted in significant losses for production observed after 72 h of exposure. These trends and the relationships between the growth parameters (Fig. 3) were similar for all materials tested, suggesting similar mechanisms for adaptation to the test concentrations of suspended matter.

Low concentrations of MP might also stimulate algal growth when the particles are both smaller (Zhao et al., 2019) and bigger (Chae et al., 2019) than cell size. In line with that, some field studies also suggest that particulate matter, under certain conditions and at concentrations <100 mg/L, stimulates algal growth instead of retarding it (Birkett et al., 2007). In our study, the algal growth was promoted by kaolin and cellulose at 10 to 100 mg/L, but not by the microplastic. At 10 mg/L, the production was significantly higher in the treatments with the reference materials than in the algae exposed to the weathered microplastic, whereas no difference from the control was observed for any of the treatment. The difference in AUC between the reference materials and PET_w_ was due to the stimulatory effect on the algal production in the kaolin and cellulose treatments. These stimulatory effects decreased in the 100 mg/L treatments and became inhibitory at 1000 mg/L in all treatments. It is unlikely that kaolin and cellulose powder provided any additional nutrients to the algae; instead, the algal growth was most probably promoted by the topography of the heteroaggregates formed by algae with kaolin and cellulose at low concentrations of suspended solids (Fig. X).

The particle aggregation occurred during the exposure in both particle controls (no algae) and all treatments with algae. Moreover, the aggregation was facilitated by the presence of algae compared to the respective particle controls, indicating that heteroaggregates, i.e., those formed by test particles and algal cells, had different PSD characteristics compared to homoaggregates, i.e., those formed by test particles only. In addition, the production of extracellular polymeric substances (EPS) by algae was a likely driver facilitating heteroaggregate formation (Costa et al., 2018). It has been suggested that EPS production is enhanced in algal cells exposed to foreign matter, such as polymer particles, leading to enhanced aggregation (Bergami et al., 2017b). Similarly, Lagarde and co-workers explained the observed variability in heteroaggregate formation as a result of different compositions of EPS produced by algae in response to polymer material in the exposure experiment (Lagarde et al., 2016), and certain algal species produce more hydrophobic EPS resulting in bigger aggregates (Chen et al., 2011; Long et al., 2015). Moreover, dose-response aggregation kinetics has been reported with a positive relationship between the aggregate size and particle concentration (Chen et al., 2011). Thus, the aggregate formation is species- and polymer-material-dependent (Long et al., 2017), which makes it challenging to account for it in microplastic hazard assessment, both methodologically and when interpreting the responses.

During the toxicity testing, the aggregation can effectively remove particles from the system, thus decreasing the exposure. For example, Fu and co-authors observed higher toxicity of PVC for algae at low levels (10 mg/L) compared to higher concentrations (1000 mg/L). They explained this phenomenon as a result of particle aggregation and sedimentation in the experimental system (Fu et al., 2019). To prevent sedimentation, we used a plankton wheel that kept the test particles and the algae in suspension during the exposure. However, particle aggregation implies a lower encounter rate between the test particles and algal cells and thus a lower probability of direct interactions. Also, heteroaggregates provide a different habitat for algae that are embedded in the aggregates, where the nutrient and light regimes may differ from that experienced by solitary cells, with concomitant effects on algal growth. We found that in all treatments, algal growth was affected primarily by the aggregate size and its variability, and to a lesser extent by the concentration of the suspended solids in the system (Table 4). Smaller aggregates supported higher growth, whereas higher SS concentration had inhibitory effects in all but kaolin treatments (Fig. S6).

No differences in any of the growth parameters were observed between the treatments with PET and PET_w_ (Fig. 1). However, there was a significantly higher within-treatment variability in the PET_w_ treatment at the highest concentration (Fig. 2). Moreover, there were significant differences in the PSD parameters between the weathered and virgin PET treatments following the exposure, with higher aggregation in the presence of algae for PET_w_ compared to PET. Also, the PSD parameters driving algal growth in PET_w_ treatment were more diverse and included variance and skewness as positive drivers, which indicates that variability of the aggregates becomes an important factor in governing algal growth in toxicity tests. Due to the changes in functional groups and an increase in surface area, weathered PVC has been reported to induce a stronger growth inhibition in freshwater alga *Chlorella vulgaris* (Fu et al., 2019). To date this has been the only work comparing the effect of virgin and aged MP on the ecotoxicological endpoints. Our findings suggest that significant growth predictors in PET_w_ were more similar to the cellulose treatment than to PET treatment (Fig. S6); thus, plastic-microorganism interactions change with weathering, but these changes are not necessarily lead to more significant impacts on biota.

In conclusion, the comparison among particle size distributions across the treatments showed that both suspended matter concentration and topography of the particle aggregates were significant growth predictors for all materials tested. We found no indication that PET particles, regardless of the weathering status, induced higher growth inhibition in a OECD-based test with unicellular algae compared to natural particles represented by kaolin and cellulose. At high concentrations, both natural and anthropogenic materials have similar capacity to cause adverse effects in algal growth inhibition tests, which must be taken into account in hazard assessment of plastic litter.

## Supplementary Materials

### Supplementary Materials and Methods

Text S1. Title of the first supplementary figure.

Text S2. Title of the second supplementary figure.

### Supplementary Figures

Fig. S1. Title of the first supplementary figure.

Fig. S2. Title of the second supplementary figure.

### Supplementary Tables

Table S1. Title of the first supplementary table.

Data file S1. Title of the first supplementary data file.

## Acknowledgments

Zandra Gerdes (ACES, Stockholm University) is thanked for suggestions on the experimental design.

## Author contributions

EG: study idea, conceptualization, data analaysis, writing; KE: experiment and methodology; SR: data interpretation, writing, conceptualization.

## Data and materials availability

Data are availanle within paper and Supplementary Information.

