## Supplemental materials for "Algal growth at environmentally relevant concentrations of suspended solids: implications for microplastic hazard assessment"

### Supplementary Material

#### Supplementary Text S1: Particle preparation and size distribution measurements

##### Test microplastic

Polyethylene terephthalate (PET; Goodfellow GmbH, product number ES306312) was obtained as 3–5 mm-sized pellets from the manufacturer and milled to a powder by Messer Group GmbH, Germany. The PET powder mixed with Milli-Q water containing 0.01 % v/v of a non-ionic surfactant (Tween 80, Sigma-Aldrich) was size-fractionated by sequential wet-sieved (200-, 100- and 40-µm sieves). Using a metal Büchner funnel with a 0.2-µm nylon membrane (Merck Millipore, GNWP04700), the <40 µm fraction was obtained; the filtrate was dried, weighed and re-suspended for size distribution analysis and further use in the experiments.

##### Reference materials

Kaolin (Sigma-Aldrich, K7375), particles of laminar shape, and native fibrous cellulose (Macherey-Nagel, MN 301) were used as the reference materials. Kaolin contains mainly the clay mineral kaolinite, a hydrous aluminosilicate, whereas cellulose is the main substance in the walls of plant cells. Both materials occur globally in suspended particulates and have been used as reference material when assessing microplastic effects (Gerdes et al. 2019) and as a test material when assessing the effects of total suspended solids (Gordon and Palmer 2015). In this work, a measured dosage of kaolin and cellulose was first added into the water of 0.5 L to prepare suspension with 10 g/L solid concentration. Then, the suspensions were mixed with an electronic stirrer for 30 min. After that, the suspensions were processed by ultrasonic disperser for 5 min to completely wet the solid particles and ensure full dispersion.

##### Particle size measurements

The particle size distribution (PSD) is a useful description of the relationship between particle abundance and size. Laser diffraction measurements are commonly used to measure and characterize suspended sediments, microorganisms and floc-size both *in situ* and in laboratory, because the measurement process is less disturbing to the flocs and aggregates compared to other methods (Serra et al. 2001). However, studies comparing particle size distributions obtained by static light scattering (volume-based estimates; vol%) and by sedimentation methods (weight-based estimates; wt%) demonstrate that static light scattering yields larger particle radii compared to sedimentation methods for identical suspensions. A limit of <8 µm (vol%) obtained by static light scattering corresponds to a limit of <2 µm (wt%) for sedimentation methods (Konert and Vandenberghe 1997; Ramaswamy and Rao 2006).

We used a laser particle counter (Spectrex, model PC-2000, Redwood City, USA) to measure PSD in all materials tested. The measurement principle is based on the light scattering by passing a rotating laser beam (wavelength 670.8 nm) through the glass container and then focusing the near-angle light pulses from the laser beam/particle collisions onto the photodetector; the collision rate is then converted into particle count and size data. Small vial attachment was used to determine the size and number of particles in the 3-100 µm range in the integrated counting mode (32 bins). The detection of particles in 2 to 3 µm range was less precise and integrated into the ≤3 µm fraction. The instrument performance was verified daily by measuring a standard of known particle size and number provided by the instrument manufacturer; as a standard, 4.2 µm polystyrene spheres in an alcohol/freon liquid matrix were used. Appropriate blanks (standard blank sample provided by the manufacturer and the particle-free water used to dilute the samples) and controls (reference samples provided by the manufacturer) were used to monitor the performance. Testing replicate samples of the polymer and kaolin standards showed that the between-replicate variation of particle counts was less than 6 % for <10 µm range and less than 3 % for < 60 µm range.

#### Particle size distribution analysis

Data on the particle size distribution obtained with the particle counter were processed using the GRADISTAT program, version 8.0 (Blott and Pye 2001) and are given according to the method by Folk and Ward (Folk and Ward 1957) to obtain mean and median ($D_{10}$) particle sizes, mode particle size for non-unimodal distributions, particle size which 10 % of the sample is below, known as $D_{10}$, particle size which 90 % of the sample is above, known as $D_{90}$, sample sorting representing variance (σ), sample skewness, and kurtosis. Sample sorting describes the spread of the particle sizes around the average particle size, where well sorted samples have low sorting values due to a low spread of the particle sizes around the average. Sample skewness describes the symmetry or preferential spread to one side of the average, where a log-normally distributed sample has a skewness value of 0. Sample kurtosis describes the degree of concentration of the grains relative to the average where a log normally distributed sample has a kurtosis value of 3. Values higher than 3 indicate a leptokurtic distribution, which when presented graphically appears strongly peaked, and smaller values indicate a platykurtic distribution, which appears relatively flat when presented graphically (Blott and Pye 2001). By using a range of statistical analyses, the linkages between algal growth, amount of suspended solids and relative abundance of microplastic in these solids on the particle distribution characteristics and aggregation can be examined.

#### Particle size distribution in the test and reference materials

The stocks of kaolin and test microplastic had comparable PSD characteristics, whereas cellulose had greater particle size range and variability (Table S1; Fig. S1). Mean particle size varied from 9.1 µm (kaolin) to 18.7 µm (cellulose), whereas these values for microplastic were intermediate and highly similar between PET and $PET_w$ (11.3 and 13.5 µm, respectively). However, the weathered plastic had substantially greater contribution of larger particles as evidenced by the D-values; in particular, the particle size which 90 % of the sample is below, known as $D_{90}$ (µm) for $PET_w$ was twice higher than for PET (Table S1). As no additional sieving was applied when producing $PET_w$ from PET, the observed increase in the mass proportion of the larger particles indicates possible aggregation of these microplastic particles in the stock mixtures that was facilitated by weathering.

#### Supplementary Figures and Tables for PSD analysis in stock suspensions

**Supplementary Table S1.** Particle size distribution parameters calculated by GRADISTAT for stock suspensions of the reference materials (kaolin and cellulose) and test microplastics, virgin and weathered polyethylene terephthalate (PET and $PET_w$, respectively); see Figure S1. Three technical replicates were obtained for each spectrum and averaged.

| **Distribution parameters, µm** | **Reference materials** | | **Microplastic** | |
|---|---|---|---|---|
| | **Kaolin** | **Cellulose** | **PET** | **$PET_w$** |
| Sample type | Unimodal | Polymodal* | Unimodal | Unimodal |
| MODE 1 | 9.500 | 11.50 | 10.50 | 11.50 |
| MODE 2 | | 17.50 | | 17.50 |
| MODE 3 | | 41.50 | | |
| $D_{10}$ | 7.602 | 10.09 | 9.645 | 9.702 |
| $D_{50}$ (MEDIAN) | 9.090 | 16.87 | 11.23 | 11.78 |
| $D_{90}$ | 10.69 | 40.47 | 13.48 | 27.92 |
| $D_{90}$ / $D_{10}$ | 1.407 | 4.012 | 1.397 | 2.878 |
| $D_{90}$ - $D_{10}$ | 3.092 | 30.38 | 3.832 | 18.22 |
| Geometric mean ($x$) | 9.107 | 18.73 | 11.28 | 13.47 |
| Sorting (σ) | 1.136 | 1.713 | 1.246 | 1.451 |
| Skewness ($Sk$) | -0.032 | 0.240 | 0.350 | 0.600 |
| Kurtosis ($K$) | 0.886 | 0.636 | 2.524 | 1.146 |

* For polymodal distribution, the sorting, skewness and kurtosis values are unreliable.

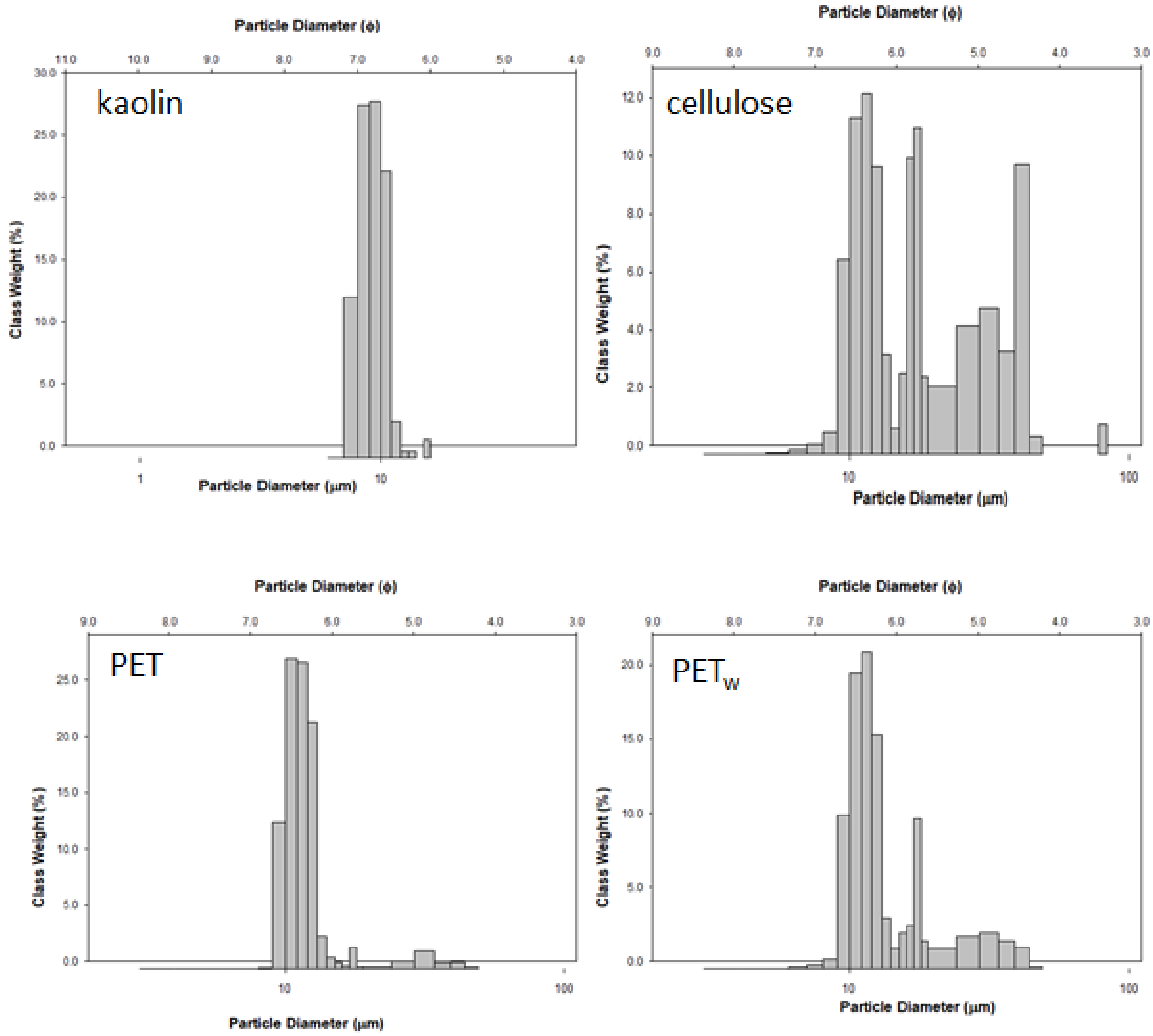


**Supplementary Figure S1.** Particle size distribution (PSD) in the stock suspensions of the reference materials (kaolin and cellulose) and test microplastics, virgin and weathered polyethylene terephthalate (PET and PETw, respectively). Output from GRADISTAT software.

**Supplementary Text S2: Algal growth model and calculation of the growth parameters**

A logistic or sigmoid curve was used to describe algal growth. When an alga is inoculated into a culture medium and exposed to suitable conditions of light, nutrients and temperature, there is a lag phase, an exponential phase, a phase of declining relative growth rate and a stationary phase. Therefore, the growth parameters considered were: lag period duration (λ) and maximum specific growth rate ($\mu_{max}$). In theory, the asymptotic value (A) will be reached once growth resources are exhausted (Fig. S1); however, in the algal growth inhibition test, this is rarely achieved.

During the lag phase, cell protein and nucleic acid contents increase and we may surmise that this phase is one of reconstitution, in which enzyme and substrate concentrations are built up to the degrees necessary for multiplication. In the exponential phase, the organisms have a high capacity for photosynthesis and the products are used mainly for the synthesis of protein. In static culture, the exponential phase ends after a time because of depletion of nutrients, accumulation of toxic by-products of metabolism, or simply because light becomes limiting as the culture becomes more dense (Fogg 1957).

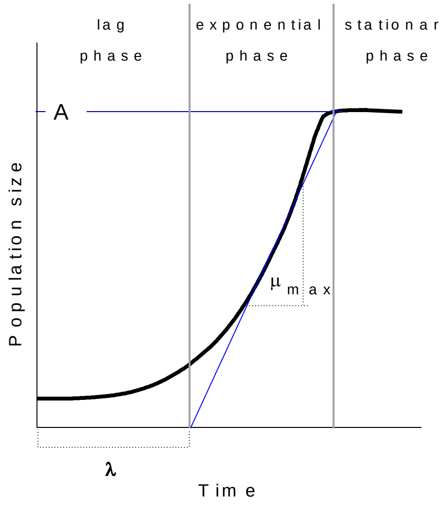


**Supplementary Figure S2.** Change in algal population size with time and the corresponding growth phases. After a lag phase (λ) characterized by zero growth, the exponential phase follows with acceleration to a maximal value ($\mu_{max}$) until the resources are exhausted resulting in ceased growth and stationary phase with zero growth.

To fit the growth profile of the algae we used a model based on a modified logistic equation (Baranyi and Roberts 1994). This model has been widely applied for various microorganisms, especially with varying lag phase (McKellar and Knight 2000; Pin et al. 2002; Baty et al. 2004; Fujikawa et al. 2004). In addition to the good fitting capacity, the model is popular because it is applicable under dynamic environmental conditions with most of the model parameters being biologically interpretable (McKellar and Knight 2000; López et al. 2004; Van Impe et al. 2005), including evaluation algal growth conditions in culture (Lacerda et al. 2011; Tevatia et al. 2012; Halmi et al. 2014; Mohamed et al. 2014). Moreover, recent evaluations suggest its superiority in growth modeling of algae under stress (Halmi et al. 2014).

The model is based on the first-order differential equation predicting specific growth rate, μ(t) ($h^{-1}$) of cell population with time (Baranyi et al., 1993; Baranyi, 1997):

$$\mu(t) = \frac{1}{X(t)}\frac{dX}{dt} = \mu_{\max}\alpha'(t)f(t) \quad \text{(Eq. 1)}$$

where $X$(t) is the algal concentration in the medium at time $t$ (expressed as total fluorescence at constant culture volume), $\mu_{max}$ is the maximum specific growth rate ($d^{-1}$), $\alpha'(t)$ is the adjustment function, and $f(t)$ is the inhibition function. In Eq. 1, $\alpha'(t)$ is a monotonous function, being $0 \leq \alpha'(t) \leq 1$ and $lim_{t\to\infty}\alpha'(t) = 1$ represents adaptation of algae in the new environment of the test conditions, i.e. the lag phase. It is based on a kinetic assumption that the growth in the lag phase is inhibited by a limiting intracellular substance following a Michaelis–Menten principle (Baranyi and Roberts, 1994; Baranyi, 1997) and expressed as:

$$\alpha'(t) = \frac{e^{-h_0}}{e^{-\mu_{max}t} + e^{-h_0} - e^{-\mu_{max}t-h_0}} \quad \text{(Eq. 2)}$$

where $h_0$ is the dimensionless model parameter (Perni et al., 2005). Finally, $f$(t) in the Eq. (1) is a monotonously decreasing function with $f(0) = 1$ and $lim_{t\to\infty}\alpha'(t) = 1$ described by the following logistic function:

$$f(t) = 1 - \frac{X}{X_{max}} \quad \text{(Eq. 3)}$$

where $X_{max}$ is maximum algal concentration observed in during the exposure and represented by the total fluorescence at constant test volume.

After rearranging Eqs. (1–3), the rate of microalgae growth ($dX/dt$) becomes:

$$\frac{dX}{dt} = \mu_{max}\left(\frac{e^{-h_0}}{e^{-\mu_{max}t} + e^{-h_0} - e^{-\mu_{max}t-h_0}}\right)\left(1 - \frac{X}{X_{max}}\right) \quad \text{(Eq. 4)}$$

The cell population in a batch reactor at time $t$ is modeled by integrating Eq. (4) at initial conditions: $X(0) = X0$ and $X(t) = X(t)$:

$$X(t) = \frac{X_{max}X_0(e^{h_0} + e^{\mu_{max}t} - 1)}{X_{max}e^{h_0} + X_0(e^{\mu_{max}t} - 1)} \quad \text{(Eq. 5)}$$

Individual (i.e., replicate-specific) growth curves were produced to derive $\mu_{max}$ and λ values. For calculations, DMFit software (www.combase.cc) was used applying models with no asymptote. None of the metapopulations have reached the stationary phase during the exposure.

#### Supplementary Text S3: PLSR modeling: details on methods and procedures

Partial least square regression (PLSR) (JMP®, Version 14.0. SAS Institute Inc., Cary, NC, 1989-2019) was used for assessing the principal latent variables (factors) underlying growth patterns in the algae exposed to the suspensions of the reference and test materials. PLSR is a bi-linear regression-modelling tool based on latent variables used to study the relation between two data tables X and Y (Martens and Martens 2001). Systematic covariations from the two data tables are decomposed into latent variables called principal components (PC). The latent variables decomposed from the X data table are in turn used for modelling the Y-variables. The model contained growth parameters ($\mu$, λ and AUC values) as response variables (Ys), whereas the candidate predictors included exposure conditions parameters: SS concentration, mean and median ($D_{50}$) particle sizes, $D_{10}$, $D_{90}$, sample variance ($\sigma$), skewness and kurtosis.

The analysis was carried out for each material as well as for the pooled data set using the non-linear iterative partial least squares (NIPALS) algorithm, with the original data matrices deconstructed into a sum of vector products for each principal component. The data were centered and scaled. In NIPALs, the scores and loadings for the first principal component are determined and then the vector product is subtracted from the initial matrix to find the residual, or unfit data. The next component is then found by analyzing the residual in the same manner.

Loadings were also calculated and plotted to give another way to view the relationships between the X- and the Y-variables and the PLS components. The loadings are determined by the independent and dependent variables, and correspond to the direction of the component in space. A larger loading for variable 1 versus 2 indicates that 1 contributes more strongly to that principal component than 2. The scores vector relates how far each observation projects along that principal component. It is important to note that even though the NIPALs algorithm is similar for PCA, the actual components differ. The PCA of the PSD characteristics will assign the highest loading values to variables that have the highest variation, while in PLSR, the emphasis on co-variation can change this order.

Because components are defined sequentially, it is important to determine how many components are needed for the optimal model. For cross-validation and determination of the optimal number of latent variables, we applied leave-one-out method and predicted residual error sum of squares (Root Mean PRESS) statistic as implemented in the PLS platform of the software. Three metrics were used to determine the optimum number of components: $R^2X$ (explains the variance) and $R^2Y$ (analogous to the coefficient of determination, $R^2$, used in regression analysis) are the coefficients of determination for the X and Y matrices, and $Q^2$ is a measure of the predictive ability of the model based on cross-validation. The cumulative value for each metric is evaluated after a component is added to the model. For each next component, the values for $R^2X$, $R^2Y$, and $Q^2$ are compared to the respective values for the previous component to determine how much including an additional component improved the fit or predictive ability or both. Because PLSR is primarily used to relate the effects of changes in X on Y, the decision to add a new component is prioritized towards the impact on $Q^2$. If $Q^2$ increases significantly, the component is retained. The algorithm will then continue until $Q^2$ either does not increase significantly or decreases. An ecological PLS model is of good quality if $R^2Y \geq 0.7$ and the $Q^2 \geq 0.4$ (Lundstedt et al. 1998)

The variable VIP (Variable Importance in Projection; a weighted sum of squares of the PLS weights) value measures explicative power of predictor variables with respect to the response variable in the PLSR. Wold and others (Wold et al. 2001) advocated cut-off-values for VIP to separate terms that do not make important contribution to the dimensionality reduction involved in PLSR (VIP $< 0.8$) and those that might (VIP $\geq 0.8$). In addition, the percentage of variation explained for X variables and Y responses and the contribution of each of the important factors were assessed. For each X variable, VIP value was calculated to assess its importance in the determination of the PLS projection model for both predictors and responses; the VIP threshold of $\geq 0.8$ was applied. Also, Xs coefficients in the PLSR model were calculated to assess their contribution to the prediction of the Ys. Based on PLSR model the predicted values for the responses were calculated and plotted versus the observed values. After preliminary runs, the important predictors were identified and included in the final PLSR prediction model (Cox and Gaudard 2013).

#### Supplementary Figures and Tables for multivariate analysis of PSD data

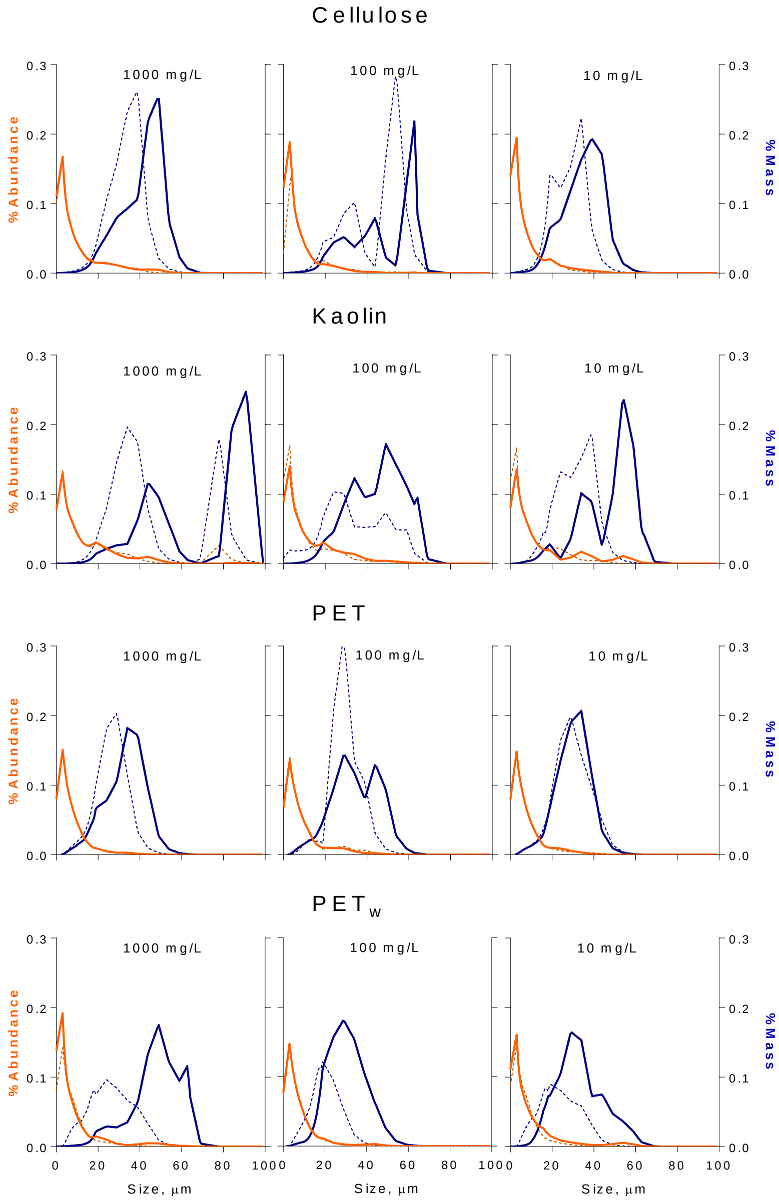


**Supplementary Figure S3.** Particle size distribution (PSD) in the experimental systems with algae (solid lines) and in the particle controls (dotted lines); see Table 1 for details on the experimental treatments. The distributions for particle number (*%Abundance*: proportion by number shown by orange line; left Y-axis) and particle mass (*%Mass*: proportion by mass, blue line; right Y-axis) are shown as average treatment values (treatments with algae: $n$ = 5, and particle controls: $n$ = 3) in the size range 3 – 100 µm. Concentration (mg/L) of the suspended solids added to the system is shown on each panel.

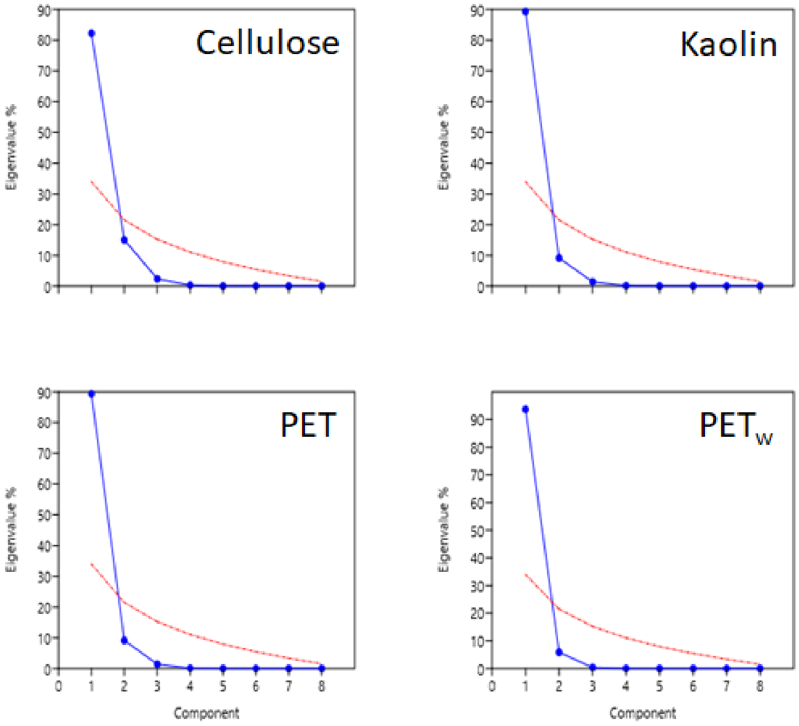


**Supplementary Figure S4.** Eigenvalues of components (blue points and the connecting line) and broken stick model (red line) identifying significant principal components for material-specific PCA models for treatments with the reference materials (kaolin and cellulose) and test microplastics, virgin and weathered polyethylene terephthalate (PET and PETw, respectively).

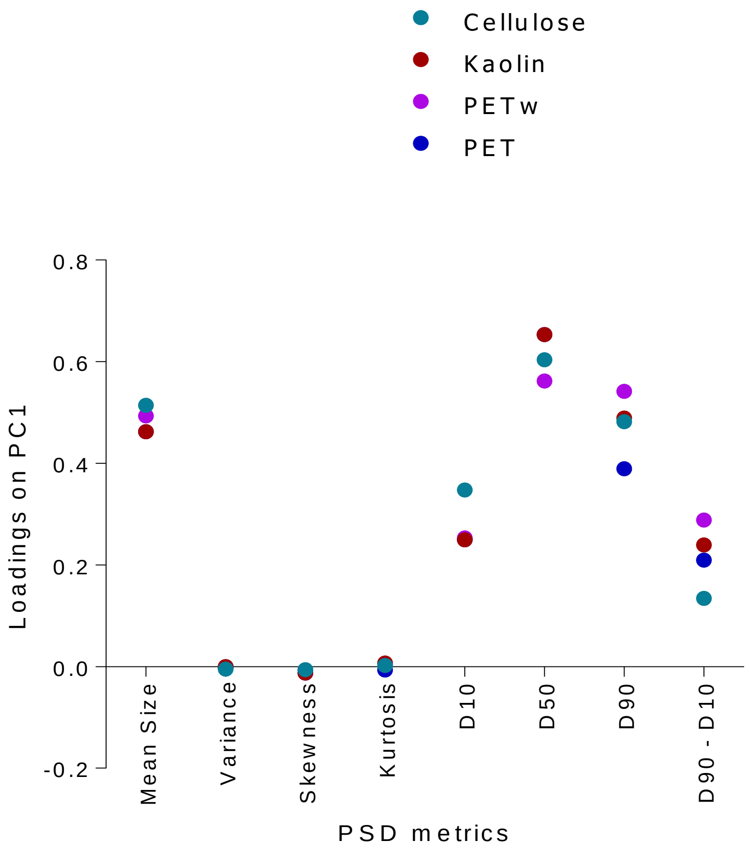


**Supplementary Figure S5.** Contribution of the data attributes to the principal components according to their loadings on PC1 for PSD in the treatments with reference materials (kaolin and cellulose) and test microplastics, virgin and weathered polyethylene terephthalate (PET and $PET_w$, respectively)

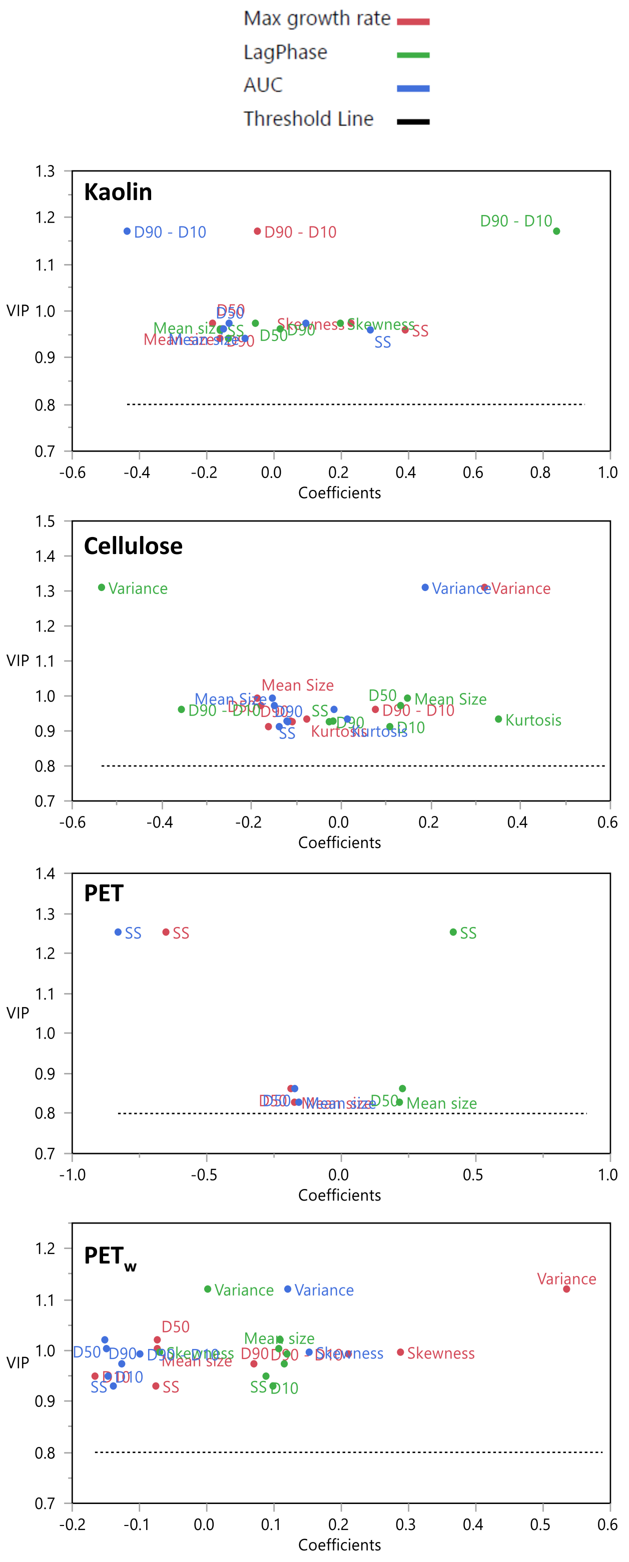


**Supplementary Figure S6.** PLSR output: VIP values vs. coefficients for independent variables in the models presented in Table 4.

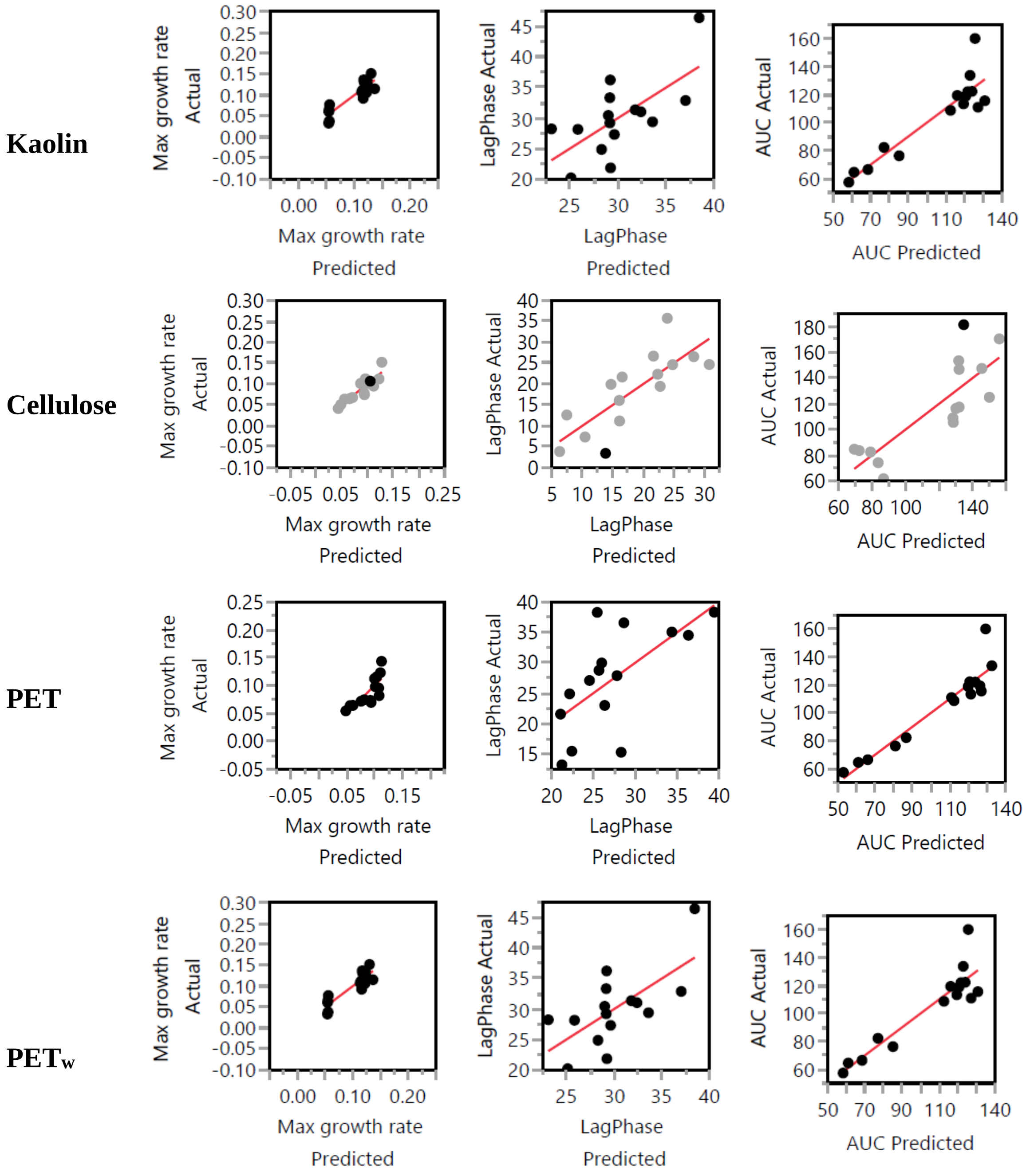


**Supplementary Figure S7.** PLSR output: Actual by Predicted plots for the dependent variables in the models presented in Table 4.
